# Brain-derived neurotrophic factor and TrkB levels in mice that lack vesicular zinc: Effects of age and sex

**DOI:** 10.1101/421453

**Authors:** Brendan B. McAllister, Nicoline Bihelek, Richelle M. Mychasiuk, Richard H. Dyck

**Affiliations:** Department of Psychology, University of Calgary, 2500 University Drive NW, Calgary, Alberta, Canada, T2N 1N4; Hotchkiss Brain Institute, University of Calgary, 3330 Hospital Drive NW, Calgary, Alberta, Canada, T2N 4N1; Alberta Children’s Hospital Research Institute, University of Calgary, 3330 Hospital Drive NW, Calgary, Alberta, Canada, T2N 4N1

**Keywords:** ZnT3, zinc transporter, synaptic zinc, SLC30A3, neurotrophin

## Abstract

In certain neurons, zinc ions are stored in synaptic vesicles by zinc transporter 3 (ZnT3). Vesicular zinc can then be released synaptically to modulate myriad targets. *In vitro* evidence indicates that these targets may include brain-derived neurotrophic factor (BDNF) and its receptor, tropomyosin receptor kinase B (TrkB). But the effects of vesicular zinc on BDNF and TrkB in the intact brain are unclear. Studies of mice that lack ZnT3 – and, as a result, vesicular zinc – have shown abnormalities in BDNF and TrkB levels, but results have been mixed and are therefore difficult to interpret. This might be caused by differences in the age or sex of mice tested. In the present study, we measured BDNF and TrkB levels in the hippocampus and neocortex, comparing wild type and ZnT3 knockout mice of both sexes at two ages (5 and 12 weeks). We also examined BDNF mRNA expression and protein levels at an intermediate age (8-10 weeks). We found that, regardless of age or sex, BDNF and TrkB protein levels did not differ between wild type and ZnT3 knockout mice. There were sex-specific differences in BDNF protein and mRNA expression, however. BDNF protein levels increased with age in female mice but not in males. And in females, but not males, ZnT3 KO mice exhibited great hippocampal BDNF mRNA expression than wild type mice. We conclude that, at least in naïve mice housed under standard laboratory conditions, elimination of vesicular zinc does not affect BDNF or TrkB protein levels.

## INTRODUCTION

The divalent cation zinc has essential biological functions throughout the body, including in the brain. Though most zinc in the brain is tightly-bound in protein structures, a portion exists in a “free” (unbound or loosely-bound) state, making it available to participate in signaling functions (McAllister & Dyck, 2017). The concentration of extracellular free zinc is relatively low, in the nanomolar range (Frederickson et al., 2006a; Anderson et al., 2015), and the cytosolic concentration is even lower, in the picomolar range (Colvin et al., 2010). However, in certain regions of the brain and spinal cord, a considerable pool of free zinc is stored in the synaptic vesicles of neurons (Pérez-Clausell & Danscher, 1985). This vesicular zinc can be released in an activity-dependent manner (Assaf & Chung, 1984; Howell et al., 1984; Aniksztejn et al., 1987), elevating the extracellular free zinc concentration to – by most estimates – the low micromolar range (Frederickson et al., 2006b), though hundreds of micromolar might be achievable with intense stimulation.

One intriguing target of zinc is brain-derived neurotrophic factor (BDNF), a member of the neurotrophin family (Barde et al., 1982). Like other excreted peptides, BDNF is produced in the cell soma as a larger precursor protein. Pre-proBDNF is cleaved intracellularly into proBDNF (32 kDa). ProBDNF is then cleaved, by furin or other proprotein convertases, to produce mature BDNF (14 kDa), which forms an active homodimer (Mowla et al., 2001). BDNF is anterogradely transported to axon terminals in dense core vesicles (Conner et al., 1997; Fawcett et al., 1997; Michael et al., 1997; Dieni et al., 2012), from where it can be released along with the cleaved pro-domain (Kohara et al., 2001; Matsumoto et al., 2008). Uncleaved proBDNF can also be released and processed extracellularly, by plasmin or matrix metalloproteinases (MMPs), into mature BDNF (Lee et al., 2001; Gray & Ellis, 2008; Nagappan et al., 2009; but see Matsumoto et al., 2008). In addition to pre-synaptic release, BDNF is also stored post-synaptically in dendrites and spines, and it can be released to act as a retrograde or autocrine signal (Wong et al., 2015; Harward et al., 2016; Choo et al., 2017).

Once released, BDNF exerts effects through tropomyosin receptor kinase B (TrkB), allosterically dimerizing these receptors and inducing their kinase function (Klein et al., 1991). This activates several signaling cascades, including the Ras, Rac, PI3-kinase, and PLC-•1 pathways (Reichardt, 2006). Truncated variants of TrkB are also expressed, with unique cytoplasmic domains that lack catalytic kinase function (Klein et al., 1990), though the T1 variant has its own BDNF-dependent signaling pathway that results in intracellular calcium release (Rose et al., 2003). When expressed in the same cells, truncated TrkB forms heterodimers with full-length TrkB and inhibits its function (Eide et al., 1996).

There is *in vitro* evidence that zinc interacts with BDNF at multiple levels. Some experiments indicate that zinc can act directly on the BDNF protein (Ross et al., 1997; Post et al., 2008; Travaglia et al., 2013). Other experiments have shown that application of micromolar concentrations of zinc to cortical cell cultures boosts BDNF messenger RNA (mRNA) expression (Hwang et al., 2011), and increases the extracellular concentration of BDNF by activating MMPs that cleave BDNF from its precursor, promoting downstream effects of BDNF (Hwang et al., 2005; Hwang et al., 2011; Poddar et al., 2016). There is even evidence that zinc can transactivate TrkB through BDNF-independent mechanisms (Huang et al., 2008; Huang & McNamara, 2012). However, uncertainty remains about how vesicular zinc interacts with BDNF in the intact brain. One way of studying this is to examine how BDNF and TrkB levels are affected in mice that lack the dedicated vesicular zinc transporter, zinc transporter 3 (ZnT3) (Palmiter et al., 1996; Wenzel et al., 1997). Elimination of this protein results in a total loss of vesicular zinc throughout the central nervous system (CNS) (Cole et al., 1999; Linkous et al., 2008). In terms of total zinc content, this corresponds with a reduction of about 20% in the cortex and 20-40% in the hippocampus (Cole et al., 1999; Lee et al., 2002; Adlard et al., 2010).

To date, several studies have examined BDNF or TrkB levels in ZnT3 KO mice, but they have produced mixed results. Adlard et al. (2010) found that hippocampal proBDNF levels, but not BDNF or TrkB levels, are reduced in ZnT3 knockout (KO) mice at 3 months of age. By 6 months, both proBDNF and TrkB levels are reduced. Similarly, Nakashima et al. (2011) showed that TrkB mRNA levels in barrel cortex are reduced in 2-month-old male ZnT3 KO mice. On the other hand, Helgager et al. (2014) showed that ZnT3 KO mice, aged 3-6 months, have normal levels of hippocampal TrkB but elevated BDNF, resulting in increased TrkB phosphorylation. And Yoo et al. (2016) found that 5-week-old male ZnT3 KO mice have increased levels of TrkB in the hippocampus and cortex, increased mature BDNF and proBDNF in the cortex, and increased mature BDNF but decreased proBDNF in the hippocampus. One obstacle to synthesizing these results is the differences in the age and sex of the mice examined, as well as differences in the brain regions assessed. Our goal in the present experiment was to address this limitation by comparing wild type (WT) and ZnT3 KO mice of both sexes at two different ages within the same study, examining BDNF and TrkB levels in both the hippocampus and neocortex. We found that neither age nor sex accounted for the disparity in past results; rather, our results suggest that elimination of vesicular zinc has no effect on BDNF or TrkB protein levels in either brain region, regardless of the age or sex of the mice tested. We did observe, however, that BDNF mRNA expression was enhanced in female, but not male, ZnT3 KO mice.

## EXPERIMENTAL PROCEDURES

### Animals

All protocols were approved by the Life and Environmental Sciences Animal Care Committee at the University of Calgary and followed the guidelines for the ethical use of animals provided by the Canadian Council on Animal Care. All efforts were made to minimize animal suffering. Mice were housed in temperature- and humidity-controlled rooms maintained on a 12:12 light/dark cycle (lights on during the day). Food and water were provided *ad libitum*. WT and ZnT3 KO mice, on a mixed C57BL/6×129Sv genetic background, were bred from heterozygous pairs. Offspring were housed with both parents until postnatal day 21, at which point they were weaned and housed in standard cages (28 × 17.5 × 12 cm with bedding, nesting material, and one enrichment object) in groups of 2-5 same-sex littermates.

### Experimental design

Brain tissue was collected from 60 mice. The initial enzyme-linked immunosorbent assay (ELISA) and Western blotting experiments used 40 mice, including WT and ZnT3 KO, male and female, and young (5-week-old) and mature (12-week-old) animals, resulting in eight experimental groups (*n* = 5 for each). Additional ELISA experiments (i.e., with acid-treated samples) and quantitative reverse-transcription polymerase chain reaction (qRT-PCR) experiments used 20 mice, including male and female animals of both genotypes (8-10 weeks of age), resulting in four experimental groups (*n* = 5 for each).

### Sample collection and protein extraction

Mice were briefly anaesthetized with isoflurane and killed by decapitation. The brain was rapidly extracted, and the neocortices and hippocampi were dissected. The neocortical samples contained primarily posterior cortex, to avoid including striatum in the sample. The extracted tissue was frozen on dry ice and stored at −80 °C.

For protein analysis, tissue samples were placed in chilled RIPA buffer (Millipore-Sigma, 50 mM Tris-HCl, 150 mM NaCl, 0.25% deoxycholic acid, 1% NP-40, 1 mM EDTA; hippocampus: 200 µl; neocortex: 400 µl) containing protease and phosphatase inhibitors (Thermo Scientific Halt Protease and Phosphatase Inhibitor Cocktail, EDTA-Free) and homogenized using a bead lyser (5 min, 50 Hz). The lysates were placed on ice for 1 h and then centrifuged at 4 °C (15 min, 12,000 *g*). The supernatants were collected, protein concentrations were determined by Bradford assay (Bio-Rad), and the supernatants were stored at −20 °C until further analysis.

For certain experiments, samples were acidified during preparation to increase the detection of BDNF (Okragly & Haak-Frendscho, 1997). The acidification procedure was adapted from Helgager et al. (2014) and was performed after the samples spent 1 h on ice, but prior to centrifugation. The pH of the samples was adjusted to <3.0 using 1 M HCl, and the samples were left for 20 min at room temperature. The pH of the samples was then adjusted to ∼7.4 using 1 M NaOH.

### ELISA

BDNF protein levels were quantified using a sandwich ELISA kit (Aviscera Bioscience, SK00752-01) that has been validated and shows good selectivity for mature BDNF over proBDNF (Polacchini et al., 2015). Within each experiment, the plates used were from the same lot and were run simultaneously. Samples were assayed in duplicate (protein loaded per well: 500 µg for hippocampus, 800 µg for neocortex for non-acid treated samples; 200 µg for hippocampus, 400 µg for neocortex for acid treated samples) following the manufacturer’s instructions. Briefly, samples were incubated for 2 h in a 96-well plate precoated with a monoclonal antibody against BDNF. The plate was then incubated for 2 h with the biotinylated detection antibody, followed by a 1 h incubation with streptavidin-HRP conjugate. Tetramethylbenzidine (TMB) substrate solution was added, and colour was developed for 10-18 min (depending on the experiment) before addition of the stop solution (0.5 M HCl). All incubations were conducted at room temperature on an orbital shaker. Absorbance was measured at 450 nm using a microplate reader (Wallac 1420 Victor^2^, Perkin Elmer Life Sciences; or FilterMax F5 Multi-Mode Microplate Reader, Molecular Devices, depending on the experiment). BDNF concentrations were determined based on a standard curve and converted to pg of BDNF per mg of total protein.

### RNA isolation and qRT-PCR

Total RNA was isolated from frozen tissue using the Allprep RNA/DNA Mini kit (Qiagen), as per the manufacturer’s protocol. Total RNA concentration and purity was determined using a Nanodrop 2000 (Thermo Fisher Scientific). Purified total RNA (2 µg) was reverse transcribed to cDNA using oligo(dT)_20_ of the Superscript III First-Strand Synthesis Supermix kit (Invitrogen), as per the manufacturer’s protocols. All primers for the target gene and reference genes were purchased from IDT (Coralville, IA). Primers were designed by an in-house technician using Primer3 (http://bioinfo.ut.ee/primer3/) to span exon-exon junctions, whenever possible.

To assess gene expression, 10 ng of cDNA with 0.5 µM of each of the forward and reverse primers for the target or reference genes and 1× PowerUP SYBR green (Thermofisher) was subjected to qRT-PCR using a Quantstudio 3 Real Time PCR system. A standard curve to determine the PCR efficiency was prepared by serial dilution of cDNA from pooled control samples from 100 ng to 0.01 ng (see Appendix Table A.1). A no-template control was also subjected to qRT-PCR per gene. The thermocycling conditions were: initial denaturation at 95 °C for 2 min, followed by 40 cycles of denaturation at 95 °C for 1 s and annealing at 60 °C for 30 s, followed by a melt curve analysis. Samples were tested in triplicate. Target gene expression was normalized to the geometric mean of the expression levels of two reference genes, *Actb* and *Ywhaz*. Relative target gene expression was determined using the 2^-•• Ct^ method (Pfaffl, 2001). Gene expression values are expressed relative to the average expression level in the WT mice (males and females combined).

### Western blotting

Protein samples were heated at 95 °C for 3 min in sample buffer containing 2% 2-mercaptoethanol. Samples were separated by electrophoresis on 12% SDS-PAGE gels (30 µg of protein per lane) and transferred to PVDF membranes (Bio-Rad). The membranes were blocked for 1 h in Odyssey tris-buffered saline (TBS) blocking buffer (LI-COR Biosciences) and incubated overnight at 4 °C in Odyssey blocking buffer containing the following monoclonal primary antibodies: rabbit anti-TrkB (1:1000, Cell Signaling Technology; #4603); mouse anti-beta-actin (1:2000; Abcam; ab8224). Blots were washed 3 × 10 min in TBS containing 0.1% Tween-20 (TBST), incubated for 1 h in Odyssey blocking buffer containing the secondary antibodies (1:10,000 anti-rabbit IRDye 800; 1:15,000 anti-mouse IRDye 680; LI-COR Biosciences), and again washed 3 × 10 min in TBST. Blots were imaged using an Odyssey infrared imaging system (LI-COR Biosciences) and quantified by densitometry using ImageJ (https://imagej.nih.gov/ij/index.html). Levels of TrkB were normalized to the level of beta-actin.

### Statistical analysis

Statistical analyses were conducted using IBM SPSS Statistics (Version 24). Unless otherwise specified, data were analyzed by three-way analysis of variance (ANOVA), with sex (male vs. female), age (5 week vs. 12 week), and genotype (WT vs. ZnT3 KO) as factors. Significant interactions were followed-up with Bonferroni-corrected simple effects tests using the pooled error term, unless equality of variances could not be assumed (Levene’s test: *p* < .05), in which case non-pooled error terms were used. All ANOVA results, including non-significant interactions, are summarized in the Appendix (Table A.2).

## RESULTS

### BDNF protein levels

Levels of BDNF in hippocampal and neocortical tissue samples from WT and ZnT3 KO mice, including both male and female and young (5-week-old) and mature (12-week-old) animals, were determined by ELISA, using a kit that is selective for mature BDNF over proBDNF (Polacchini et al., 2015). In the hippocampus (Figure 1A), no difference in BDNF levels was detected between WT and ZnT3 KO mice [*F*(1,32) = 0.36, *p* = .552]. Interestingly, we did observe that age had differing effects on male and female mice [age × sex interaction: *F*(1,32) = 7.95, *p* = .008]. Specifically, BDNF levels did not differ between 5-week-old and 12-week-old males (*p* = .151), but in females, BDNF levels increased significantly with age (*p* = .034; Bonferroni-corrected). The results from the neocortex (Figure 1B) mirrored those from the hippocampus. Levels of BDNF did not differ between WT and ZnT3 KO mice [*F*(1,32) = 0.35, *p* = .557], whereas age exerted a significant effect that differed between males and females [age × sex interaction: *F*(1,32) = 6.60, *p* = .015]. BDNF levels did not differ between 5-week-old and 12-week-old males (*p* = .355) but increased significantly with age in females (*p* = .022; Bonferroni-corrected).

**Figure 1.**
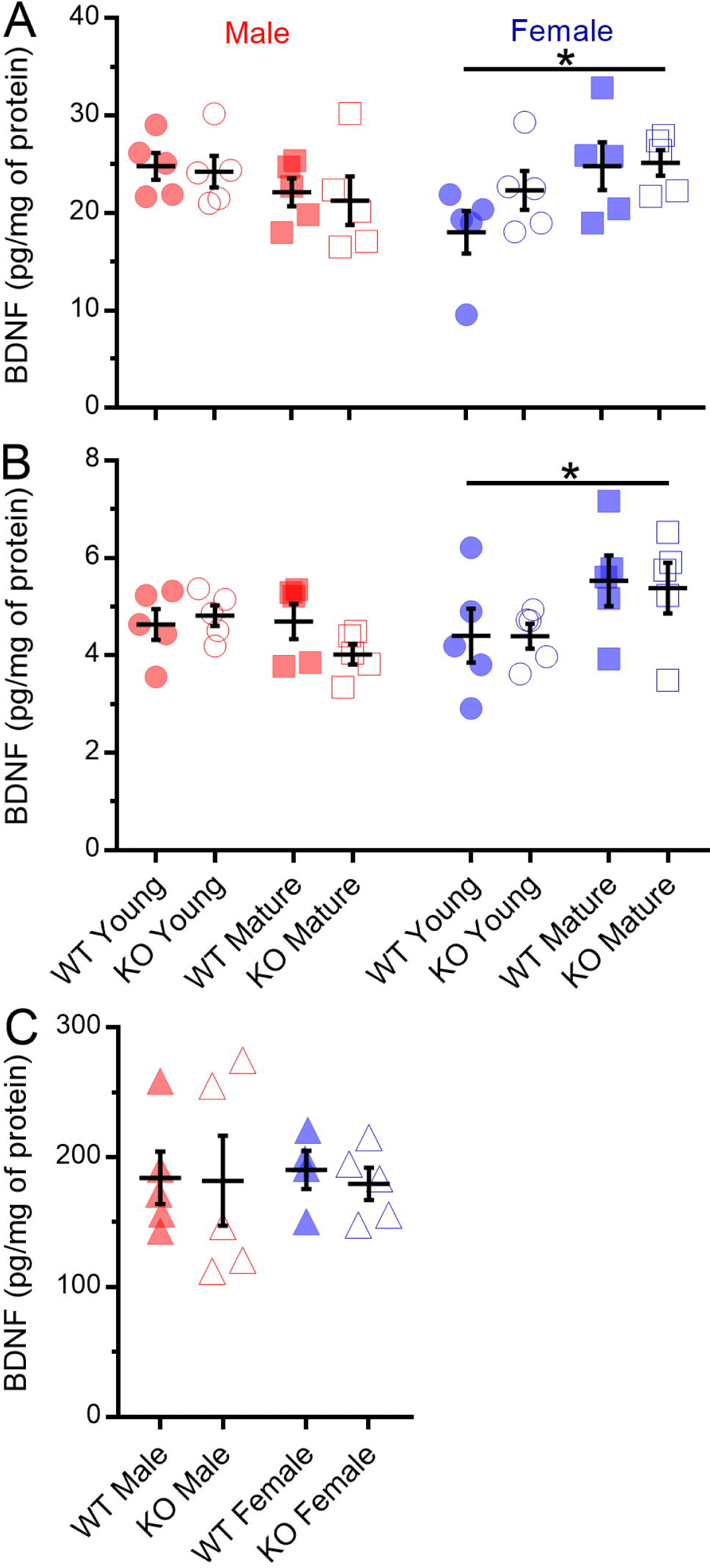
BDNF protein levels in the hippocampus and neocortex. Protein levels were quantified by ELISA. (A, B) Both hippocampal and neocortical BDNF levels were greater in mature (12-week-old) female mice than in young (5-week-old) female mice, but levels did not differ between young and mature males. There was no difference between wild type and ZnT3 KO mice. (C) Acidifying the tissue extracts during sample preparation drastically increased the detection of BDNF by the assay. However, measured BDNF levels in neocortical extracts from mice at an intermediate age (8-10 weeks) remained unaffected by genotype in both males and females. Error bars represent ±1 standard error of the mean (SEM). *follow-up test to significant interaction, *p* < .05

The BDNF levels measured by ELISA were somewhat low relative to levels described in previous reports (e.g., see Helgager et al., 2014). This could be due to methodological aspects of the sample preparation. In particular, acidification of tissue samples has been shown to increase the extraction of BDNF (Okragly & Haak-Frendscho, 1997; Kolbeck et al., 1999). Therefore, we conducted an additional ELISA experiment using hippocampal and neocortical samples that were acidified during processing. Tissue samples were taken from mice at an intermediate age (i.e., 8-10 weeks) relative to the two age groups described in the experiment above.

We confirmed that acidifying the samples dramatically increased the levels of BDNF that were detected by ELISA (compare Figure 1C to 1B). Likely, this was due to increased extraction of BDNF from the tissue, but also in part to a loss of some protein from the sample, as we noticed the formation of a considerable amount of precipitate when samples were acidified, much of which was likely lost from the sample following centrifugation and collection of the supernatant. We also noted that protein content, as determined by Bradford analysis, tended to be lower in the acidified samples than the non-acidified samples. Therefore, BDNF likely accounted for a larger proportion of the protein remaining in the sample, causing more BDNF to be detected per unit of protein loaded in the assay. By our estimate (based on total protein content of the samples), this may have accounted for ∼50% of the increase in measured BDNF concentrations. Accounting for this, the acidification procedure increased BDNF detection by approximately 20-fold, which is broadly consistent with the increased detection of other neurotrophic factors produced by acid treatment (Okragly & Haak-Frendscho, 1997).

The increase in detected BDNF concentrations was so large that, even when less protein was loaded than in our previous ELISA experiment, many of the measured values from the hippocampal samples fell outside the standard curve of the assay or outside the range of detection for the plate reader. Therefore, these data could not be used for analysis. The values measured from the neocortical samples (Figure 1C) were within the standard curve, however, and so were analyzed for differences based on the genotype and sex of the mice, using a two-way ANOVA. One sample (WT-female) was excluded from this analysis because the measured level of BDNF was very low (< 10 pg/mg of protein) relative to the other acid treated samples (range from 100-300 pg/mg of protein). Despite the improvement in BDNF detection, we were still unable to detect a difference in BDNF levels between WT and ZnT3 KO mice [*F*(1,15) = 0.08, *p* = .781]. There was also no difference between males and females [*F*(1,15) = 0.01, *p* = .937].

### BDNF mRNA levels

The results of our ELISA analysis indicated that BDNF protein levels are not abnormal in ZnT3 KO mice. We next sought to examine whether the same would apply at the level of BDNF mRNA. Brain tissue was collected from mice at an intermediate age (i.e., 8-10 weeks) relative to the 5- and 12-week-old groups described above. Samples were submitted to qRT-PCR analysis of BDNF mRNA expression. Differences based on genotype or sex were analyzed by two-way ANOVA. In the hippocampus (Figure 2A), the effect of genotype differed between sexes [genotype × sex interaction: *F*(1,16) = 5.21, *p* = .036]. Follow-up simple effects tests showed that, in females, BDNF mRNA expression was twice as great in ZnT3 KO mice than in WT mice (*p* = .002; Bonferroni-corrected), whereas in males there was no significant difference between genotypes (*p* = .372). The results exhibited a similar pattern for the neocortex (Figure 2B), although, in this case, the interaction between genotype and sex was not significant [*F*(1,16) = 1.57, *p* = .229]. The two-way ANOVA did reveal a significant main effect of genotype [*F*(1,16) = 4.90, *p* = .042], which appeared to be driven mostly by increased BDNF mRNA expression in the ZnT3 KO females relative to the WT females. However, after correcting for unequal variances, the main effect was no longer significant [*F*(1,18) = 0.65, *p* = .056; Welch’s test for unequal variances].

**Figure 2.**
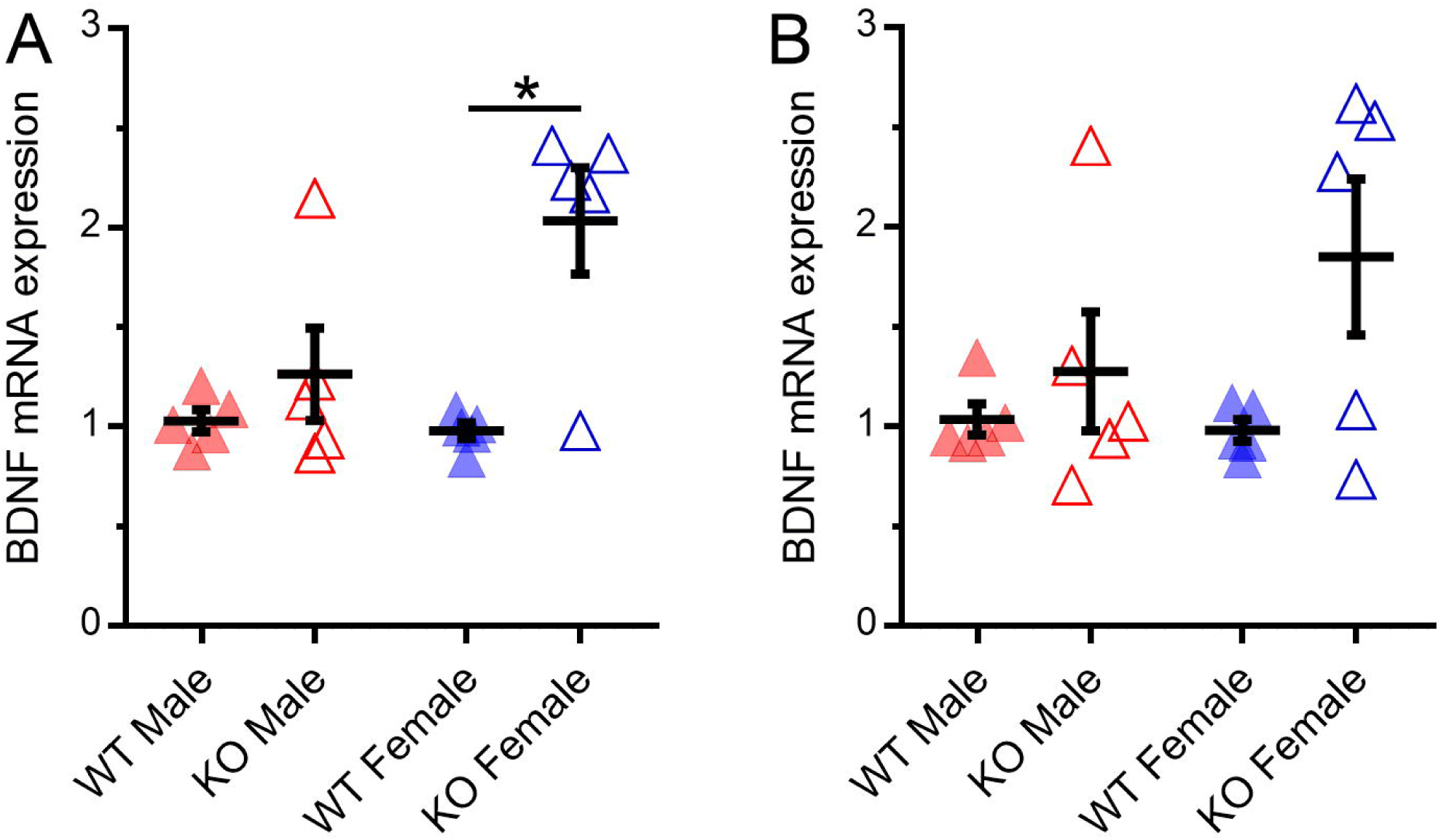
BDNF mRNA expression in the hippocampus and neocortex. mRNA expression was quantified by qRT-PCR. Target gene expression was normalized to the geometric mean of the expression levels of two reference genes, *Actb* and *Ywhaz*. Gene expression values are expressed relative to the average expression level in the WT mice (males and females combined). (A) In mice at 8-10 weeks of age, BDNF mRNA in the hippocampus was significantly greater in female ZnT3 KO mice than in female wild type mice, but did not differ between genotypes in males. (B) BDNF mRNA expression in the neocortex exhibited a similar pattern, though there was no significant effect of sex or genotype. Error bars represent ±1 SEM. *follow-up test to significant interaction, *p* < .05

### TrkB protein levels

Levels of TrkB were assessed by Western blotting (Figure 3A). The monoclonal antibody against TrkB detected two bands: one around 135 kDa, assumed to be full-length TrkB, and one around 90 kDa, assumed to be truncated TrkB (TrkB.T). TrkB.T appeared to be more abundant than the full-length version, consistent with previous findings (Fryer et al., 1996). In the hippocampus, TrkB levels (Figure 3B) did not differ between WT and ZnT3 KO mice [*F*(1,32) = 1.60, *p* = .215], nor did they differ based on sex [*F*(1,32) = 0.47, *p* = .496] or age [*F*(1,32) = 0.34, *p* = .566]. Likewise, hippocampal TrkB.T levels (Figure 3C) did not differ based on genotype [*F*(1,32) = 1.49, *p* = .231], sex [*F*(1,32) < 0.01, *p* = .958], or age [*F*(1,32) < 0.01, *p* = .959]. In contrast, TrkB levels in the neocortex (Figure 3D) were significantly higher in male mice than in female mice [*F*(1,32) = 5.13, *p* = .030]. However, there was no difference between genotypes [*F*(1,32) = 1.28, *p* = .267] or effect of age [*F*(1,32) = 0.89, *p* = .353]. Finally, levels of TrkB.T in the neocortex (Figure 3E) did not differ based on genotype [*F*(1,32) = 1.29, *p* = .265], sex [*F*(1,32) = 0.10, *p* = .757], or age [*F*(1,32) = 0.08, *p* = .777].

**Figure 3.**
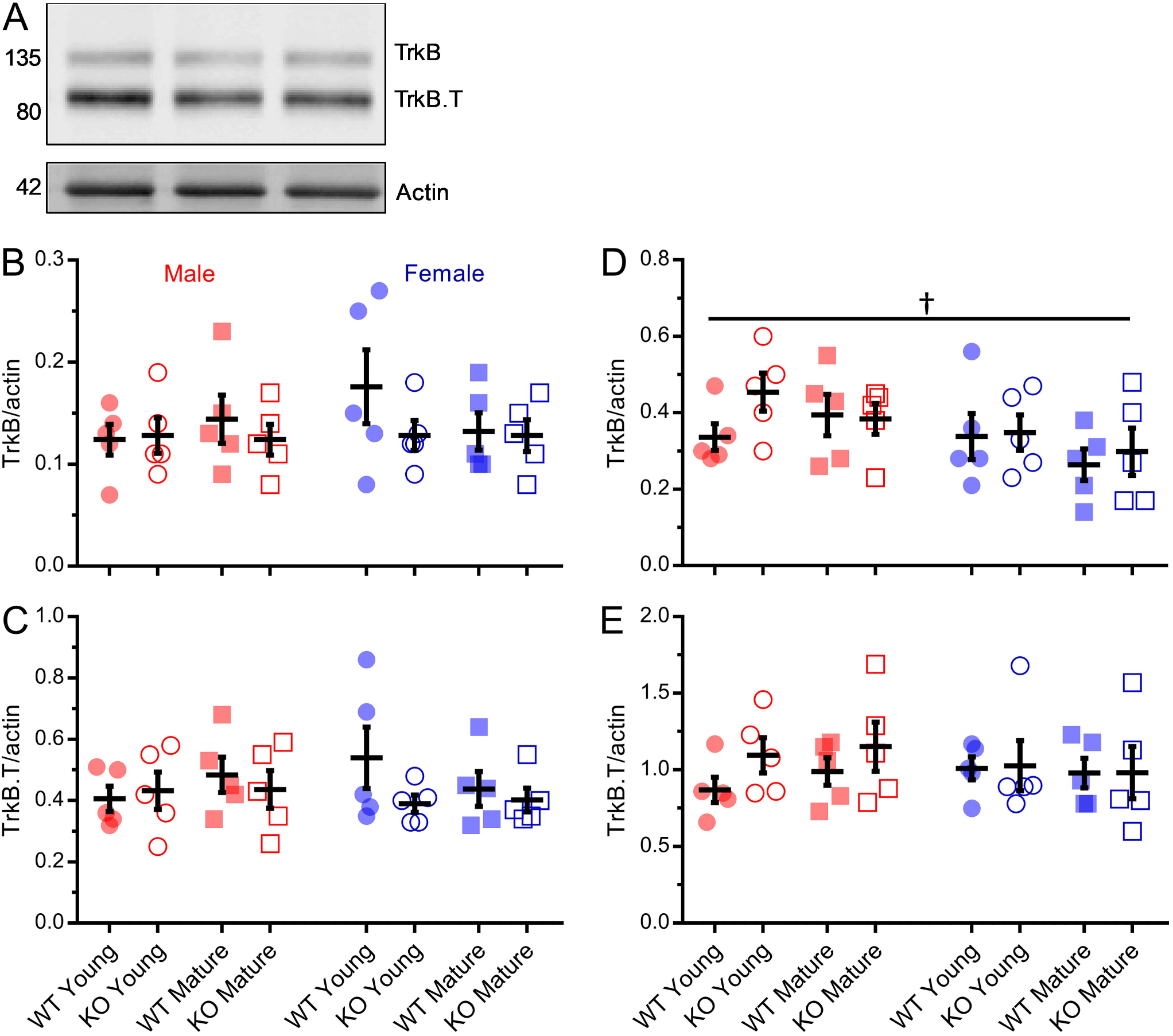
TrkB protein levels in the hippocampus and neocortex. Protein levels were quantified by densitometric measurement of Western blots. (A) Example Western blotting of hippocampal extracts (30 µg of protein loaded per lane). The antibody against TrkB detected two variants; full-length TrkB (∼135 kDa) and a truncated form of TrkB (TrkB.T; ∼90 kDa). Beta-actin was measured as a loading control, and TrkB levels were normalized to beta-actin levels. (B, C) Hippocampal TrkB and TrkB.T levels were not affected by age or sex, nor did they differ between wild type and ZnT3 KO mice. (D) Neocortical TrkB levels were greater in male mice than in female mice, but were not affected by age or genotype. (E) TrkB.T levels in the neocortex were not affected by age, sex, or genotype. Error bars represent ±1 SEM.^†^ main effect of sex, *p* < .05

## DISCUSSION

Previous examinations of BDNF levels in the brains of ZnT3 KO mice have produced seemingly discrepant results. The objective of the present study was to determine whether differences in age or sex of the mice tested might account for the discrepancy. The primary finding was that elimination of vesicular zinc, by genetic inactivation of ZnT3, did not affect hippocampal or neocortical levels of BDNF, regardless of whether mice were male or female, young or mature. Hippocampal BDNF mRNA expression, however, was increased in ZnT3 KO mice relative to WT mice, though only in females. The same appeared to apply to BDNF mRNA expression in the neocortex, though the results in this case were inconclusive. In any case, the increase in gene expression did not translate to a detectable increase in mature BDNF protein levels, though it should be noted that mRNA expression was assessed at only a single age (8-10 weeks), whereas BDNF protein levels were assessed in groups of mice at 5, 8-10, and 12 weeks of age. Finally, the absence of vesicular zinc had no effect on protein levels of the BDNF receptor, TrkB (mRNA levels for TrkB were not assessed). Thus, our study – including our analysis of age and sex – was unable to resolve the discrepancy in the literature.

Our results are in closest agreement with those of Adlard et al. (2010), who found that mature BDNF levels are normal in ZnT3 KO mice up to at least 6 months of age. They also found that hippocampal TrkB levels do not decline in ZnT3 KO mice until somewhere between 3-6 months – older than the mice examined in the present study. Previously, we found that TrkB mRNA expression in barrel cortex is reduced in male ZnT3 KO mice at 2 months of age (Nakashima et al., 2011); from the present results, it appears that this decrease in TrkB gene expression does not result in a decrease in TrkB protein, at least up to 3 months of age – though it should be noted that our analysis of TrkB protein levels extended to neocortical regions beyond barrel cortex, which could potentially mask an effect that is specific to barrel cortex. It is also possible that our Western blotting analysis may have lacked the sensitivity to detect a small change in protein levels. Our results are in partial agreement with Helgager et al. (2014), who reported that overall hippocampal TrkB levels are normal in mature ZnT3 KO mice – though TrkB phosphorylation is enhanced. However, these authors also reported that hippocampal BDNF levels are elevated, for which we found no evidence at the protein level. Our results correspond least closely with those of Yoo et al. (2016), who found elevated levels of both mature BDNF and TrkB (most likely truncated TrkB, based on their Figure 4.3B) in 5-week-old ZnT3 KO mice.

A possible explanation for the disparity in findings is the genetic background of the ZnT3 KO mice used. To our knowledge, all ZnT3 KO mice reported on to date were on a mixed C57BL/6×129Sv background strain and derived from the same original source: the laboratory of Dr. Richard Palmiter, where ZnT3 was first identified and ZnT3 KO mice were first generated (Palmiter et al., 1996; Cole et al., 1999). Our colony, initially established from mice generously provided by Dr. Palmiter, has been independently maintained over many years and generations; long enough that genetic drift may have rendered our colony a different substrain from the colonies maintained by other groups (e.g., Yoo et al., 2016) or by the Jackson Laboratory (Stock #005064), from which mice were obtained for the experiments conducted by Helgager et al. (2014). To test this possibility, the present experiments could be repeated using mice freshly obtained from the Jackson Laboratory.

One caveat to our conclusions is that our results were obtained using mice in which ZnT3 has been deleted from the germline. Thus, these mice lack ZnT3 in all cell types and throughout all of life, including development. The lack of cell specificity is not a major concern for the present experiments, but the lack of temporal specificity is an important issue, as it raises the possibility that compensatory changes could arise to counteract the effects of eliminating ZnT3, masking the effects that vesicular zinc is actually exerting in the normal brain. It is possible that more acute elimination of vesicular zinc, as could be achieved using conditional ZnT3 KO mice, would reveal different or greater effects on BDNF, as well as on the many other known or hypothesized targets of zinc. The eventual availability of such mice will greatly strengthen the field of zinc neurobiology by allowing for such possibilities to be addressed experimentally.

It is also worth highlighting that the present experiments focused only on experimentally-naïve mice. Thus, it cannot be ruled out that vesicular zinc contributes to regulating BDNF production, processing, or signaling under other conditions. To speculate, it is possible that under certain conditions wherein BDNF gene expression is enhanced – such as environmental enrichment or chronic antidepressant treatment (Nibuya et al., 1996; Tsankova et al., 2006; Zajac et al., 2010), zinc may become involved in the enzymatic processing of BDNF. It is unlikely that zinc interacts with BDNF within vesicles, as BDNF is localized to dense core vesicles (Michael et al., 1997; Dieni et al., 2012) and zinc is found in clear, round vesicles (Pérez-Clausell & Danscher, 1985). More likely is that zinc interacts with BDNF in the synaptic cleft, since both are secreted in response to neuronal activity (Frederickson et al., 2006b; Matsumoto et al., 2008; Nagappan et al., 2009; Wong et al., 2015). Assuming that enhanced BDNF expression leads to increased secretion of proBDNF, it is possible that zinc might increase the extracellular processing of proBDNF into mature BDNF by promoting the activity of MMP enzymes. Indeed, exogenous zinc application enhances MMP activity at the (probably) physiologically-relevant concentration of 10 µM (Hwang et al., 2005). Furthermore, MMP activity also appears to promote the expression of tissue plasminogen activator (Hwang et al., 2011), which, by activating plasmin, also increases the extracellular capacity to cleave proBDNF into mature BDNF (Pang et al., 2004). If vesicular zinc is indeed required for the enzymatic processing of BDNF into its mature form following increased gene expression and proBDNF production, then this would be one possible explanation for why female ZnT3 KO mice in the present study exhibited normal mature BDNF levels, despite having elevated levels of BDNF mRNA. Assuming this explanation is true, then one would hypothesize that these mice should show elevated levels of proBDNF. Unfortunately, a limitation of the present study was that proBDNF levels were not assessed; our results suggest that simultaneous measurement of pro- and mature BDNF levels in ZnT3 KO mice would be a useful undertaking in future studies.

It is also possible that, under certain conditions, zinc interacts with BDNF by influencing BDNF gene expression. Indeed, our finding of increased BDNF mRNA expression in female ZnT3 KO mice suggests that this might be the case, though it is not clear why it should apply only to females, and there is very little research on sex-differences in ZnT3 KO mice from which to formulate hypothetical explanations. Assuming that BDNF levels decline with age in ZnT3 KO mice, as suggested by Adlard et al. (2010), it is possible that increased BDNF gene expression might represent a compensatory response that is able to stave off the decline in protein levels for a period of time. It is conceivable that this age-related decline, and therefore the compensatory response, may occur earlier in females than in males. It is also worth considering possible routes through which vesicular zinc might influence BDNF gene expression. There is *in vitro* evidence that exposure to zinc can increase BDNF mRNA expression in cortical neurons, through an unknown mechanism that is dependent on MMPs (Hwang et al., 2011). Vesicular zinc release has also been shown to activate the mitogen-activated protein kinase (MAPK) pathway presynaptically in the hippocampal mossy fibers (Sindreu et al., 2011). One downstream target of MAPK is cyclic AMP response element-binding protein (CREB) (Impey et al., 1998), which can activate BDNF gene transcription (Tao et al., 2002). Another possibility is that loss of the vesicular zinc storage pool may alter intracellular zinc homeostasis; there is some evidence that the cytosolic concentration of zinc is increased in ZnT3 KO mice (Yoo et al., 2016). This requires further examination, but – if true – it would open a wide range of possible mechanisms by which zinc could interact with intracellular pathways, potentially leading to altered BDNF expression.

Though we were unable to detect any effect of eliminating ZnT3 on BDNF protein levels, we did observe some unanticipated effects of age and sex that were independent of ZnT3 genetic status. In both the hippocampus and neocortex, BDNF protein levels increased with age – from 5 weeks to 12 weeks – in female mice, but not in male mice. The topic of sex differences in BDNF levels is complex, with findings that vary between studies (Chan & Ye, 2017). For example, hippocampal BDNF levels have been found to be higher in female rats than in male rats (Bakos et al., 2009), but the opposite in mice (Szapacs et al., 2004). Regarding how BDNF levels change with age, other researchers have previously noted an increase in levels over the first few postnatal weeks in mice and rats (Katoh-Semba et al., 1997; Kolbeck et al., 1999; Kellogg et al., 2000; Silhol et al., 2005; Yang et al., 2009). It is possible that BDNF expression peaks slightly later in females, which could explain why levels continue to increase after 5 weeks, whereas in male mice they apparently do not. Also, BDNF mRNA and protein expression in the hippocampus and other brain regions can be regulated by estradiol and other sex hormones, in complex fashion (Gibbs, 1999; Jezierski & Sohrabji, 2000; Solum & Handa, 2002; Carbone & Handa, 2013), and estradiol levels do tend to rise in female mice from 5- to 12-weeks of age (Hill et al., 2012). On the other hand, Hill et al. (2012) measured hippocampal and frontal cortical mature BDNF levels over time in both male and female C57BL6 mice, and did not find that BDNF levels were clearly higher at 12 weeks than at 5 weeks in either sex, though transient sex- and region-dependent peaks were observed prior to 12 weeks.

We also observed that full-length (but not truncated) TrkB levels were higher in male mice than in female mice, though only in the neocortex. The cause of this difference is not clear. Hill et al. (2012) provide some evidence that BDNF and TrkB levels tend to be inversely regulated over time; thus, if BDNF levels increase with age in females, as our data suggest, one might expect TrkB levels to be lower in females. However, hippocampal and frontal cortical TrkB levels have been directly compared between male and female mice at 2.5-4 months of age in other studies, and no sex differences were found (Hill et al., 2011, 2013). A limitation of the present study is that mRNA expression data were not collected to expand on the findings of sex- and age-dependent differences in BDNF and TrkB protein levels, as BDNF mRNA expression was not examined at multiple ages, and TrkB mRNA expression was not assessed at all.

The primary conclusion of the present study is that – contrary to previous reports (Helgager et al., 2014; Yoo et al., 2016) – BDNF protein levels in the hippocampus and neocortex are not affected by the absence of vesicular zinc in naïve mice housed under standard laboratory conditions. Furthermore, we found no evidence that differences in age or sex of the mice tested could explain discrepant findings in the literature on BDNF levels in ZnT3 KO mice; this discrepancy is likely the result, therefore, of other unidentified methodological factors. Further research will be required to clarify the conditions under which ZnT3 and vesicular zinc interact with BDNF processing or signaling, with a particular focus on how the effects of vesicular zinc differ at the level of BDNF gene expression and enzymatic processing, and the mechanisms by which these effects might be influenced by sex.

## ACKNOWLEDGEMENTS

The authors thank Dr. Frank Visser and the HBI Molecular Core facility for use of their equipment and for consultation, as well as Dr. Tuan Trang for use of his laboratory’s plate reader and for helpful suggestions. The authors also thank Dr. Visser and Melinda Wang for technical assistance with the qRT-PCR analysis.

## APPENDIX

**Table A.1.**
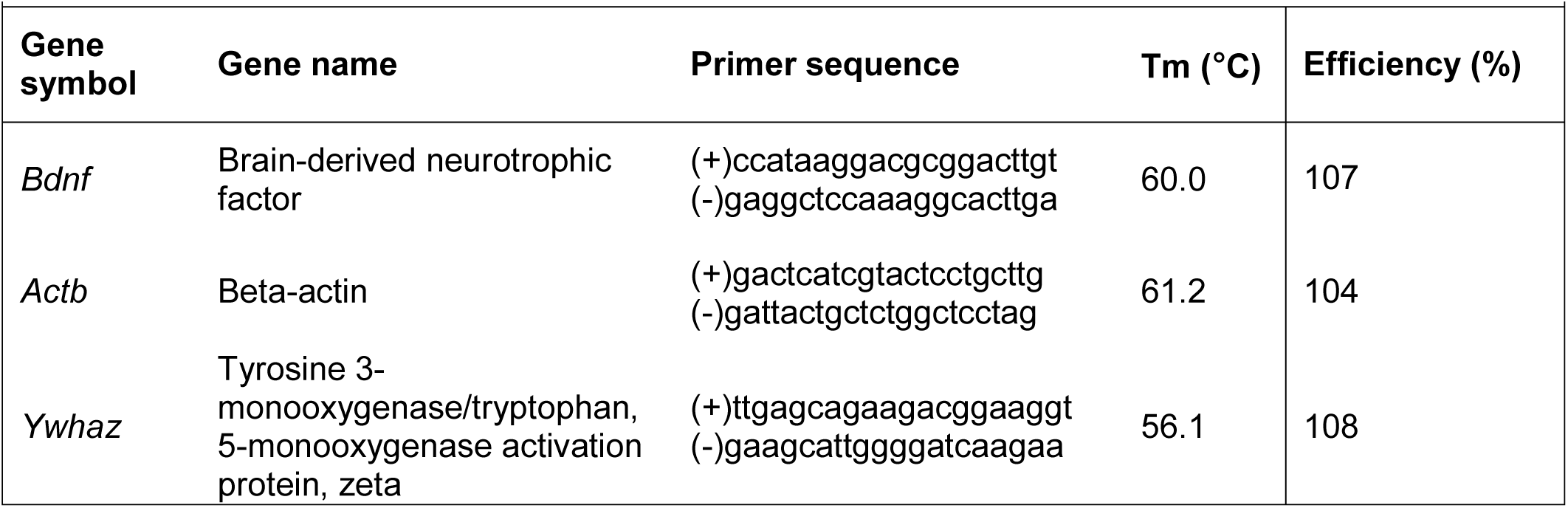
Supplemental information for qRT-PCR methods. Primer sequences and melting temperatures for qRT-PCR conducted to assess mRNA expression of BDNF and two reference genes.

**Table A.2.** ANOVA results. Main effects of age, sex, and ZnT3 genotype; and two- and three-way interactions between these factors. Degrees of freedom (D.F.).

|  | D.F. | Age | Sex | Genotype |
| --- | --- | --- | --- | --- |
| **Neocortex** |  |  |  |  |
| BDNF (ELISA) | 1, 32 | $F = 1.54, p = .224$ | $F = 1.92, p = .176$ | $F = 0.35, p = .557$ |
| BDNF (ELISA with acid treatment) | 1, 15 | N/A | $F = 0.01, p = .937$ | $F = 0.08, p = .781$ |
| TrkB (Western blot) | 1, 32 | $F = 0.89, p = .353$ | **$F = 5.13, p = .030$** | $F = 1.28, p = .267$ |
| TrkB.T (Western blot) | 1, 32 | $F = 0.08, p = .777$ | $F = 0.10, p = .757$ | $F = 1.29, p = .265$ |
| Actin (Western blot) | 1, 32 | $F = 1.75, p = .195$ | $F = 2.00, p = .167$ | $F < 0.01, p = .980$ |
| BDNF mRNA (normalized) | 1, 16 | N/A | $F = 1.07, p = .317$ | **$F = 4.90, p = .042$** |
| *Actb* mRNA expression | 1, 16 | N/A | $F = 0.51, p = .488$ | $F = 0.44, p = .515$ |
| *Ywhaz* mRNA expression | 1, 16 | N/A | $F = 0.02, p = .905$ | $F = 0.32, p = .577$ |
| **Hippocampus** |  |  |  |  |
| BDNF (ELISA) | 1, 32 | $F = 0.55, p = .466$ | $F = 0.15, p = .697$ | $F = 0.36, p = .552$ |
| TrkB (Western blot) | 1, 32 | $F = 0.34, p = .566$ | $F = 0.47, p = .496$ | $F = 1.60, p = .215$ |
| TrkB.T (Western blot) | 1, 32 | $F < 0.01, p = .959$ | $F < 0.01, p = .958$ | $F = 1.49, p = .231$ |
| Actin (Western blot) | 1, 32 | $F = 0.17, p = .680$ | $F = 0.03, p = .869$ | $F = 0.03, p = .860$ |
| BDNF mRNA (normalized) | 1, 16 | N/A | $F = 4.01, p = .062$ | **$F = 12.83, p = .002$** |
| *Actb* mRNA expression | 1, 16 | N/A | $F = 0.30, p = .592$ | $F = 0.23, p = .637$ |
| *Ywhaz* mRNA expression | 1, 16 | N/A | $F = 1.98, p = .178$ | $F = 0.58, p = .458$ |

**Table A.2 (Continued). ANOVA results.**
|  | Age × sex | Sex × genotype | Age × genotype | Age × sex × genotype |
| --- | --- | --- | --- | --- |
| **Neocortex** |  |  |  |  |
| BDNF (ELISA) | **$F = 6.60, p = .015$** | $F = 0.09, p = .761$ | $F = 0.80, p = .377$ | $F = 0.42, p = .523$ |
| BDNF (ELISA with acid treatment) | N/A | $F = 0.04, p = .854$ | N/A | N/A |
| TrkB (Western blot) | $F = 0.60, p = .446$ | $F = 0.21, p = .649$ | $F = 0.54, p = .470$ | $F = 1.16, p = .290$ |
| TrkB.T (Western blot) | $F = 0.52, p = .477$ | $F = 1.09, p = .304$ | $F = 0.06, p = .816$ | $F = 0.02, p = .904$ |
| Actin (Western blot) | $F = 0.08, p = .776$ | $F = 0.04, p = .840$ | $F = 0.25, p = .619$ | $F = 0.08, p = .777$ |
| BDNF mRNA (normalized) | N/A | $F = 1.57, p = .229$ | N/A | N/A |
| *Actb* mRNA expression | N/A | $F = 2.15, p = .162$ | N/A | N/A |
| *Ywhaz* mRNA expression | N/A | $F = 1.74, p = .206$ | N/A | N/A |
| **Hippocampus** |  |  |  |  |
| BDNF (ELISA) | **$F = 7.95, p = .008$** | $F = 1.26, p = .270$ | $F = 0.62, p = .438$ | $F = 0.45, p = .508$ |
| TrkB (Western blot) | $F = 1.01, p = .322$ | $F = 0.46, p = .504$ | $F = 0.07, p = .795$ | $F = 1.51, p = .228$ |
| Truncated TrkB (Western blot) | $F = 1.07, p = .309$ | $F = 0.97, p = .332$ | $F = 0.06, p = .814$ | $F = 1.22, p = .278$ |
| Actin (Western blot) | $F = 1.10, p = .302$ | $F = 0.15, p = .705$ | $F = 0.01, p = .921$ | $F = 1.35, p = .254$ |
| BDNF mRNA (normalized) | N/A | **$F = 5.21, p = .036$** | N/A | N/A |
| *Actb* mRNA expression | N/A | $F = 0.80, p = .384$ | N/A | N/A |
| *Ywhaz* mRNA expression | N/A | $F = 0.47, p = .503$ | N/A | N/A |

